# Superhelicase-driven isothermal amplification of repetitive and low-complexity DNA

**DOI:** 10.1101/2024.11.27.625726

**Authors:** Fahad Rashid, Jimin Kang, Tiantian Shang, Peter J. Murray, Xiaotian Bi, Raquel Merino-Urteaga, Momčilo Gavrilov, Wonyoung Jo, Armaan Ahmed, Qianyun Yan, Tural Aksel, Doug Barrick, James M. Berger, Taekjip Ha

## Abstract

PCR is a cornerstone of molecular biology, but many biologically important DNA templates remain difficult to amplify. Long tandem repeats, low-complexity tracts, and sequences with extreme base composition often yield low product levels, smeared bands, stutter products, or truncated amplicons. These failures can arise because repeated cycles of thermal denaturation and reannealing promote off-register annealing, polymerase slippage, secondary-structure formation, and incomplete extension. Previously, we developed SSB-Helicase Assisted Rapid PCR (SHARP), an isothermal amplification method in which an engineered superhelicase and single-stranded DNA-binding protein replace the thermal melting step of PCR with enzymatic strand separation. Here, we tested whether SHARP can improve amplification of templates that are refractory to conventional PCR. SHARP robustly amplified up to six identical tandem repeats of the Widom 601 nucleosome-positioning sequence and up to 35 identical ankyrin repeats, targets that were poorly amplified by conventional PCR under the conditions tested. SHARP also amplified templates with extreme base composition, including a 95% AT-rich template and GC-rich templates (up to ∼95% GC), as well as a 99-repeat CGG tract associated with Fragile X syndrome and a CAG/CAA repeat tract from mutant HTT exon 1 associated with Huntington’s disease. Together, these results show that helicase-driven isothermal amplification can expand access to repetitive and compositionally extreme DNA sequences that are challenging for conventional thermocycling-based PCR.

## INTRODUCTION

The Polymerase Chain Reaction (PCR) has transformed molecular biology by enabling exponential, sequence-specific amplification of DNA from small amounts of starting material^1–3^. Since its development, PCR has become central to cloning, sequencing, diagnostics, forensics, and many other applications^4–7^. Despite its broad utility, PCR has limitations when it comes to amplifying certain types of DNA, including long tandem repeats, low complexity tracts and sequences with extreme base composition^8–11^. These difficulties arise from the way PCR relies on repeated cycles of strand denaturation and re-annealing. For tandem repeats, denatured products can anneal out of register because adjacent repeat units contain similar or identical sequence. Incompletely extended products can also anneal within the repeat tract and act as aberrant primers in subsequent cycles^9,12–14^. In addition, DNA polymerases can slip during synthesis across repetitive templates, causing the nascent strand to realign incorrectly on the template. These events generate products with altered repeat number, truncated products, or mixed amplicon populations that appear as smeared or laddered bands on gels^9,12,13,15^.

These effects are compounded for templates with extreme base-composition bias. GC-rich regions form unusually stable duplexes and secondary structures (e.g., hairpins or G-quadruplexes) that resist denaturation and stall polymerases, whereas AT-rich/low-complexity regions promote mis-annealing and polymerase pausing^16–19^. As a result, incomplete extensions and template switching become more frequent, amplification is biased toward shorter variants, and a single, clean amplicon often fails to dominate.

These limitations are important because repetitive and compositionally biased sequences are widespread in biology. Repetitive DNA comprises a large fraction of eukaryotic genomes and can influence gene regulation, chromosome stability, segregation, and genome evolution^20–22^. Expanded tandem repeats are associated with multiple human disorders, including Huntington’s disease, fragile X syndrome, and myotonic dystrophy^23,24^. Repetitive sequences are also used in pathogen typing and diagnostic applications, including repetitive elements in *Mycobacterium tuberculosis*, such as MIRU-VNTR (Mycobacterial Interspersed Repetitive Unit-Variable Number Tandem Repeat) and IS6110-based approaches^25,26^. Separately, extreme base composition can complicate amplification of biologically relevant loci, such as AT-rich sequences from *Plasmodium falciparum* or GC-rich human loci such as FMR1 (Fragile X Messenger Ribonucleoprotein 1)^27–30^. Improved methods for amplifying these challenging templates would therefore be useful for both research and diagnostic workflows.

Several approaches have been developed to address these challenges, including nested PCR, touchdown PCR, hot-start PCR, specialized polymerases, additives, loop-mediated isothermal amplification, and helicase-dependent amplification. These methods can be effective in specific cases, but they often require extensive optimization, altered primer design, or template-specific reaction conditions. Consequently, sequences that are poorly amplified by standard PCR may require cloning, PCR-free cloning, synthesis-based approaches, or other labor-intensive recovery strategies. A useful alternative would preserve the simplicity and primer compatibility of PCR while reducing the failure modes caused by repeated thermal denaturation and reannealing^14,31–37^.

We previously developed SSB-Helicase Assisted Rapid PCR (SHARP), an isothermal DNA amplification method in which enzymatic strand separation replaces the thermal melting step of PCR^38^. SHARP uses an engineered PcrA superhelicase together with single-stranded DNA-binding protein (SSB), a strand-displacing DNA polymerase, ATP, and conventional PCR primers. In this reaction, the helicase unwinds duplex DNA, SSB stabilizes the exposed single-stranded templates, and the polymerase extends annealed primers under isothermal conditions. Because SHARP does not require repeated cycles of global template denaturation and reannealing, we reasoned that it could reduce off-register annealing, secondary-structure formation, and slippage-associated heterogeneity during amplification of repetitive or compositionally extreme DNA.

Here, we test SHARP on a panel of templates that frequently confound conventional PCR **(Fig. 1)**. These include tandem Widom 601 nucleosome-positioning sequence arrays, ankyrin repeat arrays, an extremely AT-rich template, GC-rich targets from plasmid and genomic DNA, and a mutant HTT exon 1 repeat-containing template. Across these examples, SHARP produced cleaner and more discrete amplicons than PCR under the conditions tested. These results show that helicase-driven isothermal amplification can expand access to repetitive and compositionally extreme DNA sequences that are challenging for conventional PCR.

**Figure 1.**
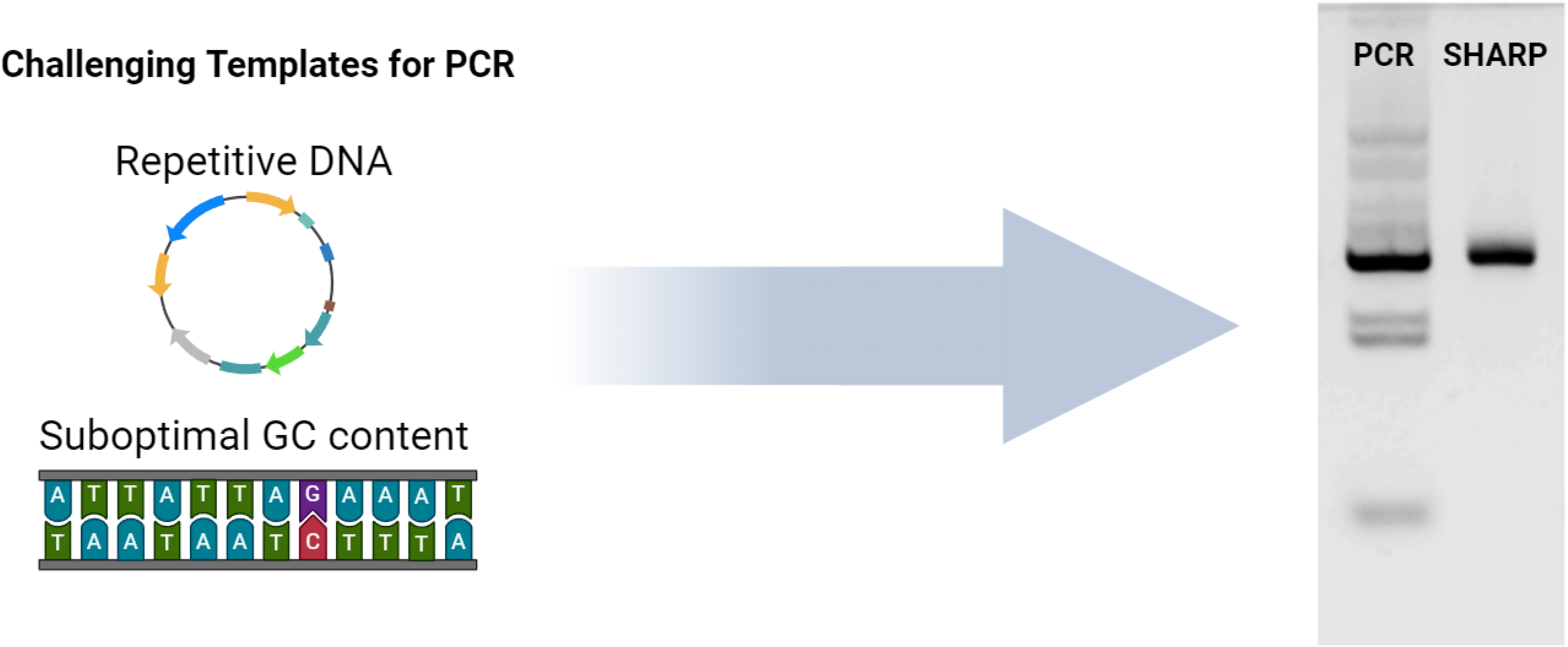
Schematic representation of the challenges faced by PCR and the solutions offered by SHARP. PCR struggles with amplifying repetitive DNA sequences and templates with suboptimal GC content, often leading to inefficient amplification. On the other hand, SHARP handles these challenging templates by employing rapid isothermal DNA amplification. SHARP’s ability to overcome the limitations of PCR makes it a powerful tool for amplifying complex nucleic acid sequences, enabling new possibilities in various fields of research and diagnostic applications.

## RESULTS

### Optimization of the crosslinked PcrA superhelicase for SHARP amplification

We previously developed SHARP using an engineered PcrA superhelicase constrained in an unwinding-competent conformation through a disulfide bond between two strategically introduced cysteins^38^. In the present study, we used BMOE to covalently crosslink the engineered cysteines, thereby generating a more uniform helicase preparation that was less sensitive to variation in cysteine oxidation state. We first examined the effects of ionic strength and pH on SHARP activity. Amplification was strongest at 50–75 mM KCl **(Fig. S1A)**, and robust activity was observed across the pH range tested **(Fig. S1B)**.

### SHARP robustly amplifies tandem repeats of nucleosome positioning sequence

We designed and cloned constructs containing multiple tandem Widom 601 sequences, a well-characterized and widely used synthetic nucleosome positioning sequence with strong affinity for histone octamers. The Widom 601 sequence we used is the symmetric 601 variant^39^. The constructs included di-, tetra-, and hexa-repeats of the Widom 601 sequence, separated by linker regions of either 65 bp or 80 bp **(Fig. 2A)**. The full sequences of the constructs can be found in the Supplementary Material.

**Figure 2.**
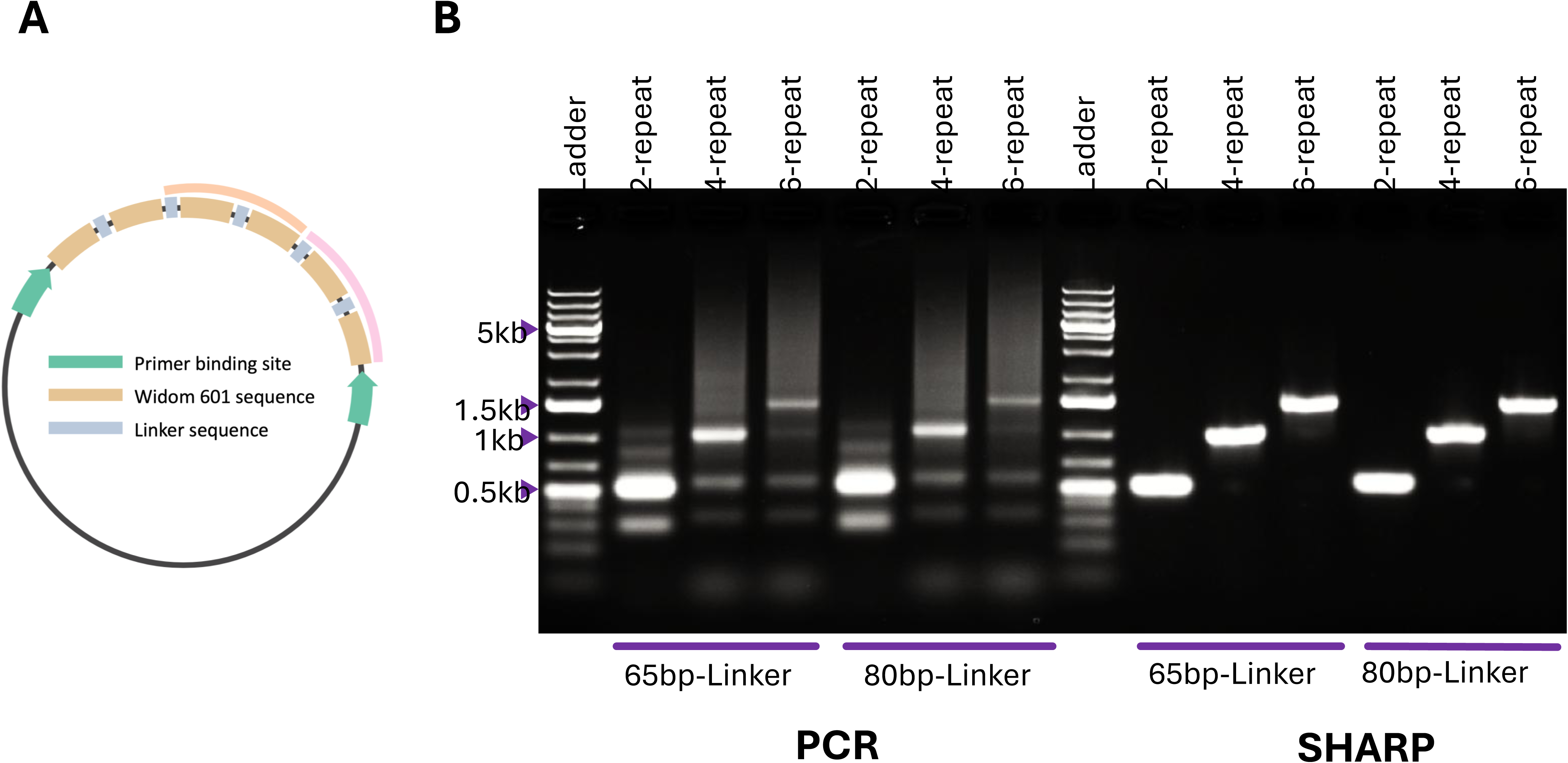
SHARP outperforms PCR in amplifying Widom 601 repeat constructs. (A) Schematic of the plasmid templates containing tandem Widom 601 repeats separated by either 65-bp or 80-bp linker sequences. **(B)** Conventional PCR amplification of constructs containing 2, 4, or 6 Widom 601 repeats with 65-bp or 80-bp linkers. PCR produced reduced yield and heterogeneous products as repeat number increased. **(C)** SHARP amplification of the same constructs generated discrete products near the expected sizes, demonstrating improved amplification of repetitive Widom 601 arrays.

Using identical primer pairs, we compared amplification of these repeat arrays by conventional PCR versus SHARP **(Fig. 2B)**. As repeat number increased, standard PCR produced progressively poorer outcomes: yields decreased and the reaction increasingly generated heterogeneous products, including smearing and off-size bands consistent with mis-annealing and slippage within the repeat tract. These artifacts were most pronounced for the longer arrays, where amplification became inefficient and poorly defined.

In contrast, SHARP generated strong, discrete products at the expected sizes for all repeat numbers tested. Notably, SHARP maintained robust amplification for the 6-repeat constructs, where conventional PCR performed worst. This improvement was observed for both linker designs (65-bp and 80-bp), indicating that SHARP performance is not restricted to a single spacer architecture. Together, these results show that SHARP substantially improved amplification of tandem Widom 601 repeat arrays relative to conventional PCR under the conditions tested, producing higher-yield and more discrete products.

### SHARP enables efficient amplification of long ankyrin repeat arrays

To test SHARP on a different class of repetitive DNA with increasing length and internal homology, we used a panel of plasmids containing tandem ankyrin repeats flanked by unique primer binding sites **(Fig. 3A)**. The repeat number spanned a wide range (1, 2, 3, 5, 9, 11, and 35 repeats), providing a stringent benchmark to test for slippage and mis-annealing prone amplification in tandem repeat sequences. We amplified each construct using the same primer pair and compared conventional PCR with SHARP **(Fig. 3B–C)**. Under standard PCR conditions, amplification quality deteriorated as repeat number increased. While the 1–3 repeat templates yielded detectable products, higher-repeat templates produced markedly reduced yield and increased heterogeneity, with poor amplification evident from 5 repeats onward and little to no robust product for the longest arrays **(Fig. 3B).** In contrast, SHARP produced strong, discrete bands at the expected sizes across the full series of repeat lengths, including the largest 35-repeat construct **(Fig. 3C)**. The SHARP reactions also exhibited a clear stepwise increase in product size with repeat number, consistent with faithful amplification of the intended repeat arrays rather than collapse or rearrangement during amplification. Together, these data show that SHARP expanded the accessible size range for tandem ankyrin repeat amplification under the conditions tested, maintaining product yield and definition as conventional PCR became increasingly inefficient and heterogeneous with increasing repeat number.

**Figure 3.**
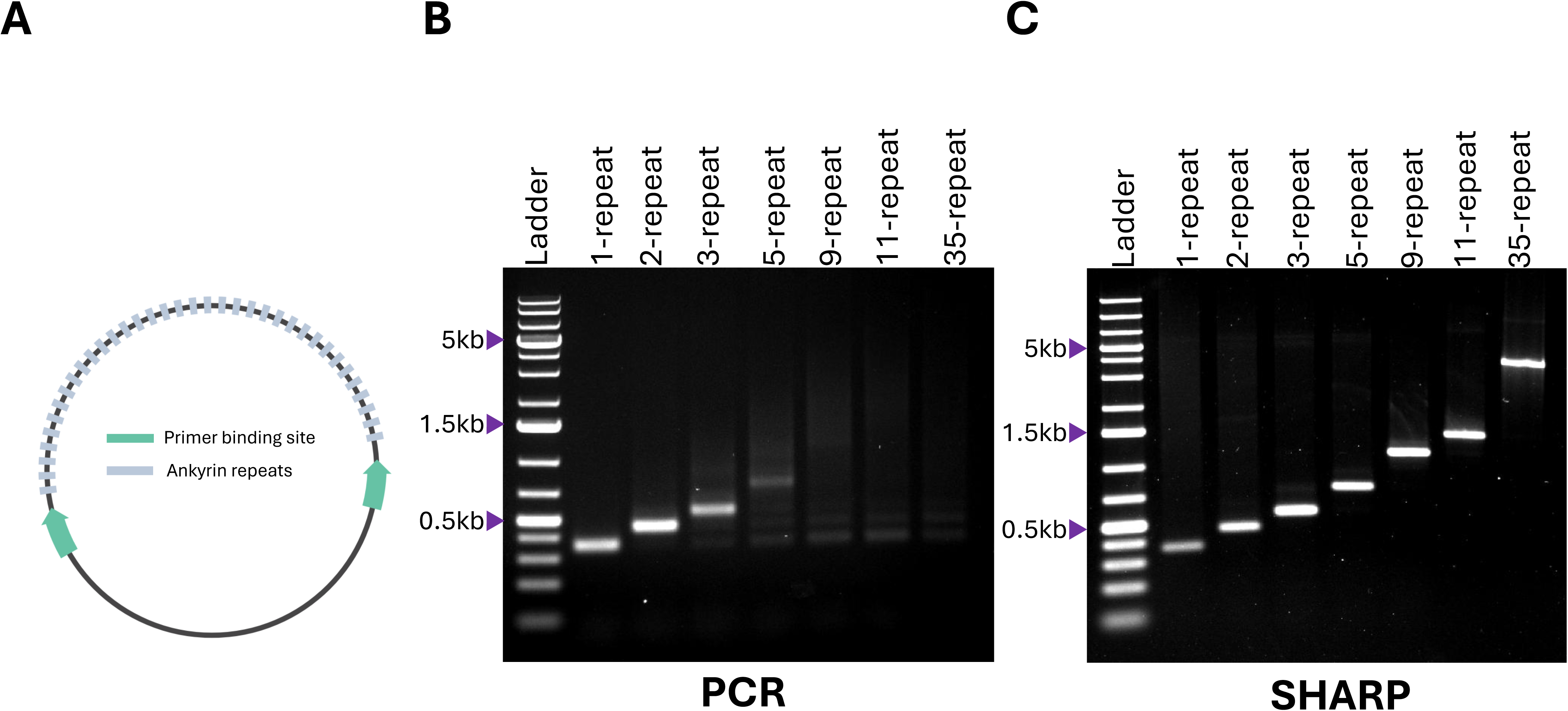
SHARP amplifies Ankyrin repeat constructs of increasing length and complexity. **(A)** Plasmid schematic of ankyrin, visualizing the context for the repetitive sequence being amplified. **(B)** PCR amplification of Ankyrin repeat constructs, showing limited efficiency from 4 repeats onward. The gel image displays that as the number of repeats increases (5,9,11 and 35), PCR struggles to amplify the sequences. **(C)** SHARP amplification of Ankyrin repeat constructs, demonstrating consistent and efficient amplification across all tested repeats (1,2,3,5,9,11,35). SHARP maintains strong band intensities, indicating superior performance in amplifying these challenging repetitive sequences.

### SHARP amplifies an extremely AT-rich template and yields a homogeneous product

Next, we tested whether SHARP could amplify DNA with extreme base composition, templates which are known to be difficult for standard PCR. We designed a 561-bp target that is 95% AT, containing extended low-complexity tracts of 235 bp and 186 bp of near-continuous A/T sequence **(Fig. 4A)**. Such templates are particularly challenging for thermal cycling–based amplification because low sequence complexity increases mis-annealing and polymerase slippage, often resulting in poor yield and heterogeneous products. Using identical primer-binding sites, we compared conventional PCR and SHARP on this AT-rich template **(Fig. 4B)**. Under the conventional PCR conditions tested, amplification was weak and poorly defined. In contrast, SHARP produced a strong, discrete band at the expected size, indicating that helicase-driven strand separation supports efficient amplification even when base composition is highly skewed toward AT. To further assess the fidelity and homogeneity of the SHARP product, we analyzed the resulting amplicons by single-molecule optical tweezers DNA unzipping **(Fig. 4C)**. Highly AT-rich sequences have been previously utilized for single-molecule DNA unzipping experiments in optical tweezers^31^. In these experiments, optical tweezers are used to mechanically pry apart double-stranded DNA, detecting the force required to unzip each base pair. Because GC base pairs interact more strongly than AT base pairs, the force-extension profile reveals unfolding transitions that reflect the underlying DNA sequence^32, 33^ **(Fig. 4C)**. AT-rich unzipping constructs reduce the background force needed to separate DNA, which for studying some systems, such as bound proteins or DNA bubbles, enables the detection of otherwise obscured, subtle features. By ligating the SHARP amplified product to DNA handles tethered between functionalized beads, we unzipped the construct and observed reproducible unfolding transitions matching those of the plasmid amplified by traditional cloning **(Fig. 4C**). In addition, the regions of relatively low and high force corresponded to the regions of minimal and moderate GC content. Taken together, these observations support the fidelity and homogeneity of the amplified product.

**Figure 4:**
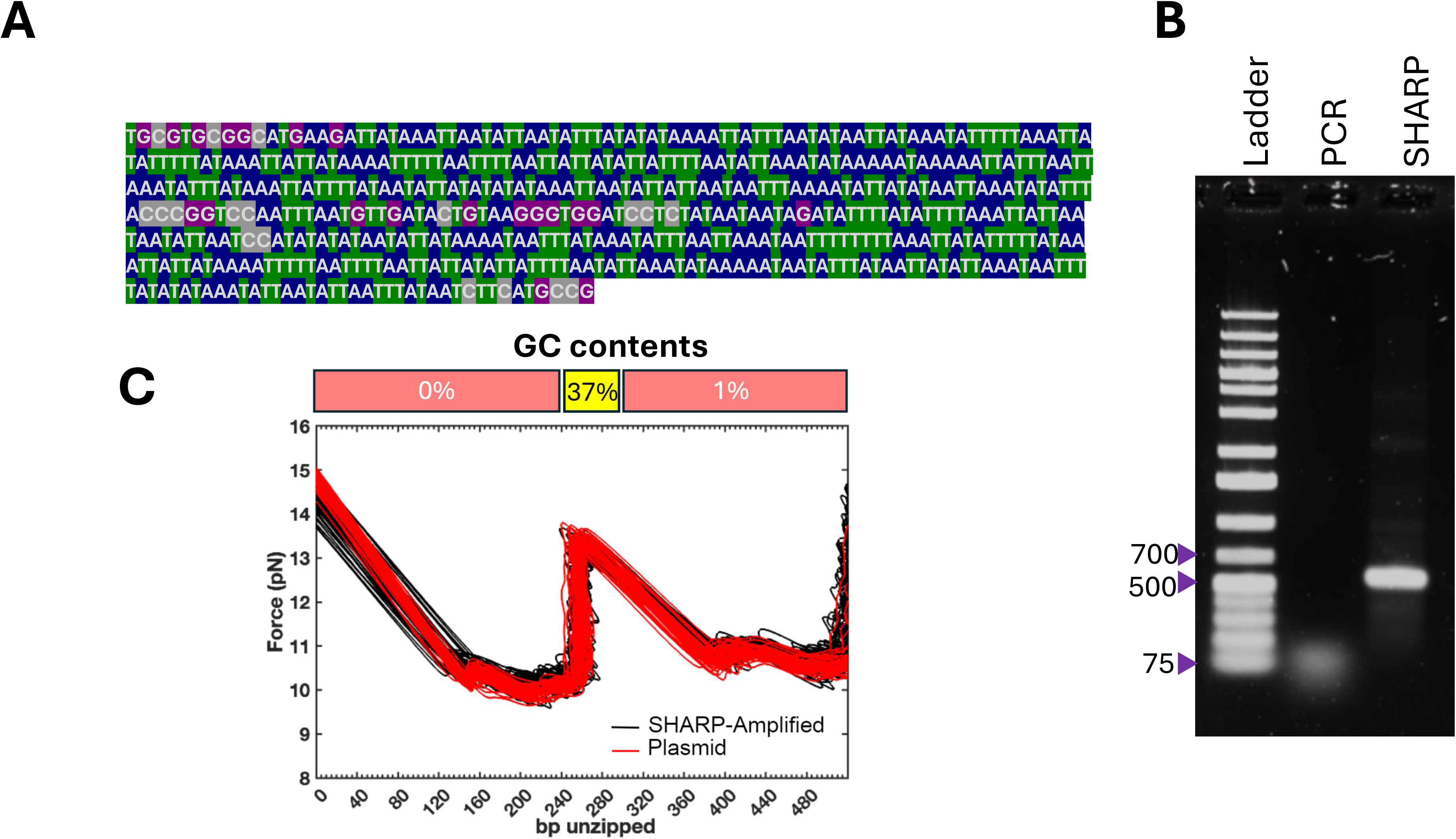
SHARP amplifies an extremely AT-rich sequence. **A)** Sequence of the AT-rich SHARP amplicon. The amplicon is 561 bp long, has 95% AT-content, and includes stretches of 235 and 186 consecutive AT base pairs. **(B)** An agarose gel comparing SHARP and PCR amplification products of the AT-rich sequence. Unlike PCR, SHARP amplification achieves a clear band at the expected length. **(C)** Optical tweezers unzipping force-extension curves validating the fidelity and homogeneity of the SHARP-generated AT-rich amplicons compared to the plasmid template (Plasmid n=40, SHARP Amplicon n=60). As expected, the regions of relatively low force correspond to the long AT stretches, while the higher-force peak in the middle corresponds to the 37% GC-content segment.

### SHARP enables amplification of extremely GC-rich targets from plasmid and genomic DNA

We next asked whether SHARP can amplify templates at the opposite extreme of base composition, where high GC content and stable secondary structures often render conventional PCR inefficient or unusable. We tested two GC-rich targets: (i) an extremely GC-rich (∼95% GC) sequence, containing 99 repeats of CGG cloned in a plasmid, and (ii) a GC-rich (∼83% GC) segment from the 5′ UTR of FMR1 amplified directly from genomic DNA **(Fig. 5A)**. These targets represent sequences that can be challenging to amplify by conventional PCR because of strong duplex stability and secondary-structure formation. Using identical primer sets for each target, we compared amplification by SHARP versus conventional PCR **(Fig. 5A)**. Under the conventional PCR conditions tested, both targets produced weak or heterogeneous products with prominent smearing, consistent with inefficient amplification of GC-rich templates. In contrast, SHARP generated clear, discrete bands for both targets from plasmid and genomic DNA, indicating efficient and specific amplification. Notably, the improvement was evident even for the genomic FMR1 target, where template complexity and potential structure formation further challenge amplification. Together, these results demonstrate that SHARP supports amplification across extreme GC content and from both plasmid and genomic DNA sources, including targets that were poorly amplified by conventional PCR under the conditions tested.

**Figure 5:**
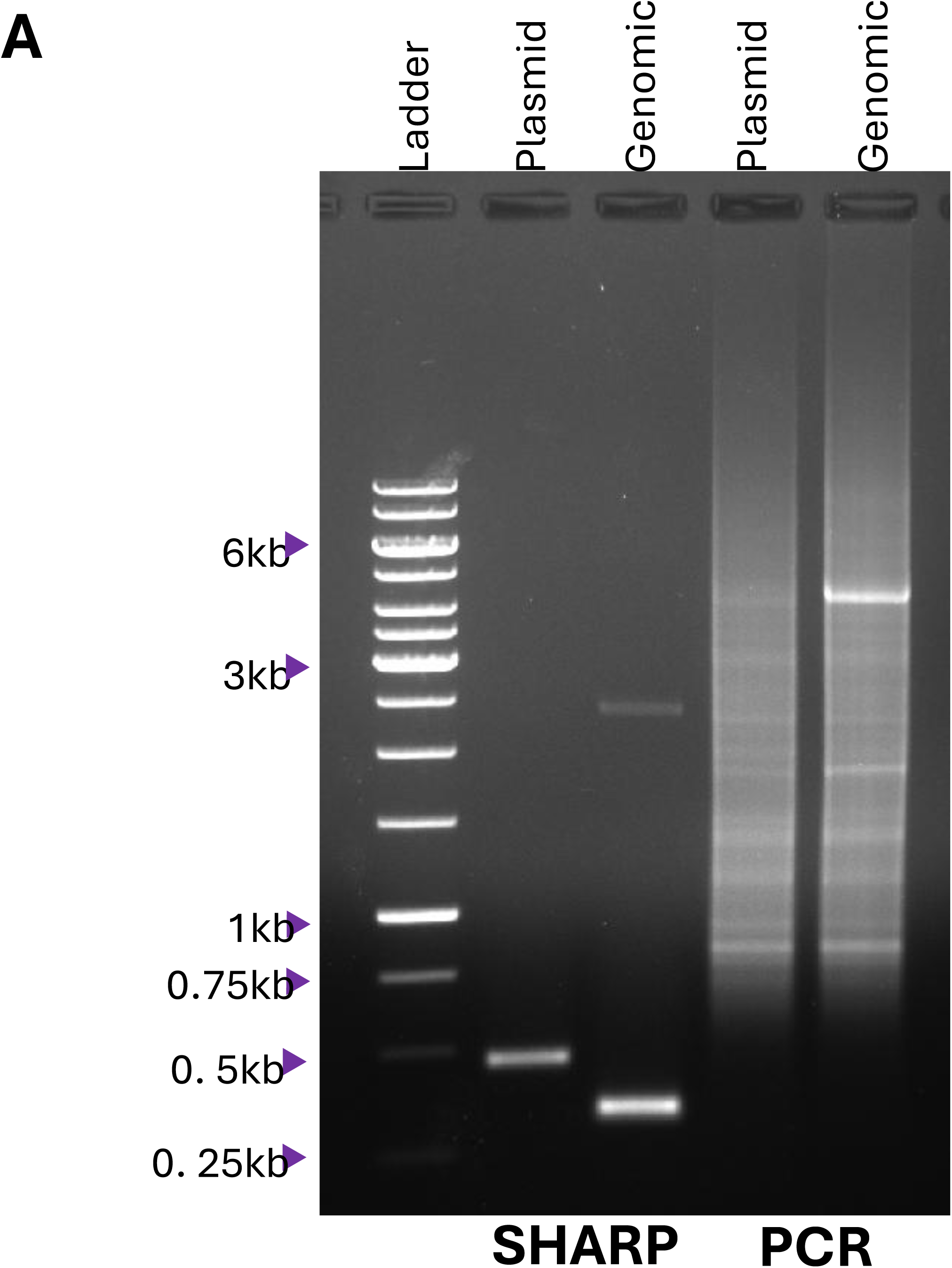
SHARP amplifies an extremely GC-rich sequence. **A)** SHARP amplifies 95% GC rich sequence from plasmid DNA as well as 83% GC content sequence of 5’UTR sequence of FMR1 gene from genomic DNA. Both targets were poorly amplified under the conventional PCR conditions tested, whereas SHARP produced discrete products at the expected sizes.

### SHARP reduces amplification heterogeneity in an mHTT repeat-containing template

Given SHARP’s demonstrated ability to accurately amplify tandem repeats, we sought to explore its suitability for repeat expansion-associated disorders such as Huntington’s diseases (HD). These conditions are characterized by long, unstable copy number of tandem repeats that are difficult to resolve via PCR due to high side-product burden and polymerase slippage. To test this, we used a plasmid template containing a mutant HTT gene encoding exon 1 of mutant Huntington protein containing a 97Q tract and benchmarked the performance of SHARP against conventional PCR **(Fig. 6)**. The tract is composed of 48 copies of CAG interspersed with 49 copies of CAA, both coding for glutamine (Q).

**Figure 6.**
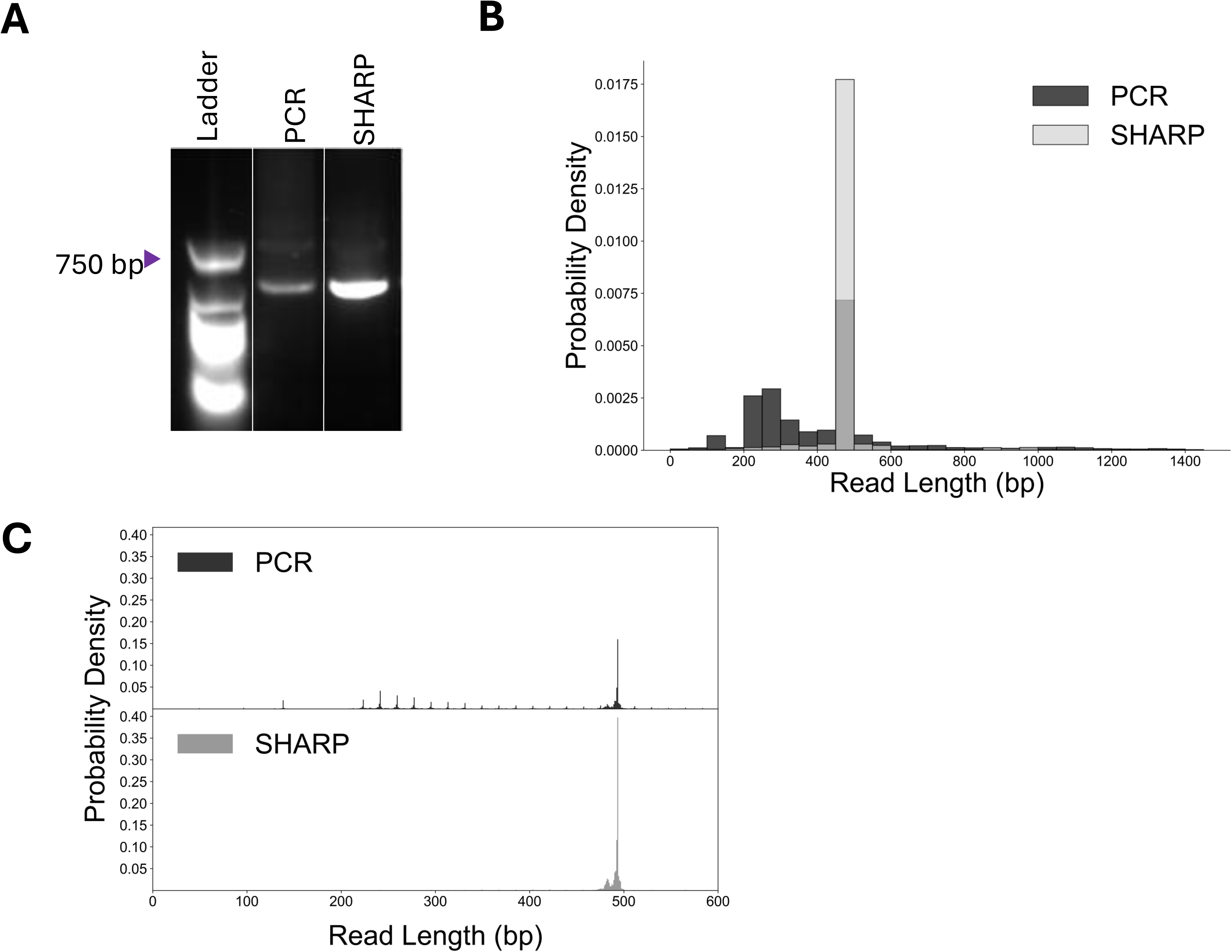
SHARP amplifies a mutant HTT gene encoding exon 1 of mutant Huntington protein containing a 97Q tract.(A) Agarose gel analysis showing amplification of the mutant HTT gene by both PCR and SHARP. 100 ng total DNA was loaded each lane. **(B, C)** Read length distributions of PCR- and SHARP-derived amplicons determined by single-molecule nanopore sequencing with 50-bp bin (B) and 1-bp bin (C). SHARP products (gray) show a predominant peak at the expected 492 bp, whereas PCR products (black) display a broader length distribution.

For the 492 bp target amplicon, equal amounts of DNA from the PCR (utilizing Q5 high-fidelity polymerase) and the SHARP reaction were analyzed. Gel electrophoresis revealed that SHARP produced a markedly more intense and discrete band at the expected size, whereas PCR yielded a characteristic smear indicating high heterogeneity **(Fig. 6A)**. To further characterize the amplicons at single-molecule resolution, we performed nanopore sequencing **(Fig. 6B, C)**. Under the conditions tested, conventional PCR exhibited a broad distribution of reads and numerous stutter peaks below the target length, whereas SHARP produced a dominant, narrow peak centered at the expected length. The read-length distribution of the PCR products exhibited periodic peaks spaced by approximately 18 bp. This spacing is consistent with the underlying (CAG)_3_(CAA)_3_ architecture of the repeat tract, in which each composite repeat block spans 18 bp. These results suggest that SHARP can reduce amplification-associated heterogeneity in repeat-containing templates. Further testing with uninterrupted CAG repeat tracts and genomic samples will be needed to evaluate its potential for analysis of disease-associated repeat expansions.

## DISCUSSION

PCR has transformed molecular biology by enabling rapid, sensitive, and sequence specific DNA amplification for cloning, sequencing, diagnostics, forensics, genetic analysis, and biotechnology. Despite this broad utility, some DNA templates remain difficult to amplify by conventional thermocycling-based PCR. Long tandem repeats, low-complexity tracts, and sequences with extreme AT or GC content remain challenging for conventional PCR to amplify.

Here, we tested whether SHARP could improve amplification of difficult templates. SHARP replaces the thermal melting step of PCR with enzymatic strand separation by an engineered PcrA superhelicase, while SSB stabilizes exposed single-stranded DNA and a strand-displacing polymerase extends conventional PCR primers. We propose that local helicase-driven strand separation reduces opportunities for off-register reannealing and secondary-structure formation compared with repeated global denaturation and reannealing during PCR **(Fig. 7)**.

**Figure 7.**
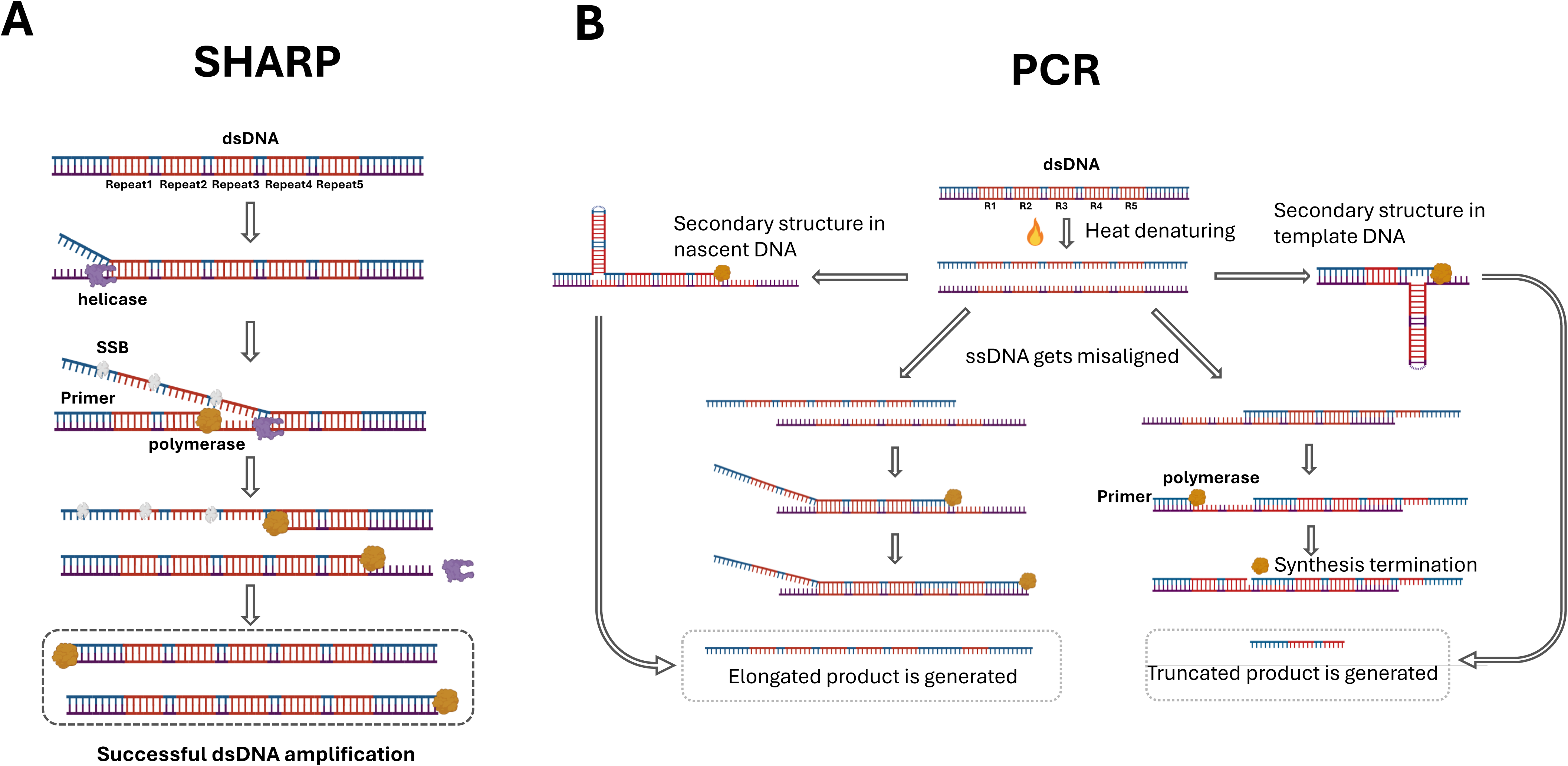
Schematic comparison between SHARP and PCR for repetitive sequence amplification. **(A)** SHARP pathway showing steps of helicase-mediated unwinding, SSB binding, primer annealing, and DNA synthesis for successful amplification. **(B)** PCR amplification showing how heat denaturation can cause elongated or truncated products through misalignment and secondary structure formation of repetitive sequences.

Consistent with this model, SHARP robustly amplified several classes of repetitive DNA that were poorly handled by conventional PCR. Tandem Widom 601 nucleosome positioning arrays and ankyrin repeat arrays became increasingly difficult to amplify by PCR as repeat number increased, generating weak, smeared, or heterogeneous products. In contrast, SHARP produced stronger and more discrete amplicons at the expected sizes across the repeat series, including templates that were inefficiently amplified or not detectably amplified by PCR. SHARP also improved amplification of a mutant HTT exon 1 repeat-containing template. For this target, conventional PCR produced a heterogeneous smear and broad nanopore read-length distribution, whereas SHARP generated a cleaner gel product and a narrower read-length distribution centered near the expected amplicon size. These findings suggest that SHARP can reduce repeat-associated amplification artifacts, although additional benchmarking will be needed to fully define repeat length accuracy and sequence fidelity.

SHARP also performed well on templates with extreme base composition. Under the conditions tested, conventional PCR did not generate discrete products from the 95% AT-rich template or the highly GC-rich targets, whereas SHARP generated strong, discrete products. In contrast, SHARP generated strong and discrete products from these templates. These results suggest that helicase-driven isothermal amplification can help overcome two distinct sources of PCR failure: mis-annealing and low sequence complexity in AT-rich regions, and high duplex stability or secondary-structure formation in GC-rich regions.

We compared SHARP with a standardized conventional PCR workflow under the conditions tested but did not independently optimize specialty-PCR conditions for each target using alternative polymerases, additives, primer designs, or cycling programs. Therefore, some templates that were poorly amplified by conventional PCR under these conditions may be recoverable through target-specific PCR optimization. Further improvements in SHARP may also come from optimizing helicase, polymerase, and SSB combinations or engineering enzymes with higher processivity, stability, and specificity.

Although SHARP successfully amplified a plasmid-borne CGG repeat tract and the mHTT repeat-containing template, direct amplification of long repetitive expansions from complex human genomic DNA remains challenging in the current iteration of the assay. Amplification of repetitive tracts from plasmids is inherently less demanding than from genomic DNA, where a vast excess of non-target sequence is present. We hypothesize that in complex genomic backgrounds, this excess DNA acts as a competitive sink for SSB and PcrA, reducing their effective concentrations at the target locus and hindering processive unwinding. Importantly, SHARP amplification from genomic DNA was demonstrated in our original report, and in the present study, SHARP successfully amplified a GC-rich, non-repeat segment of the FMR1 5′ UTR directly from genomic DNA. This result indicates that genomic DNA complexity *per se* is not the primary barrier; rather, the combination of repeat-tract secondary structure combined with a complex genomic background appears to be the limiting factor. As a more immediate strategy, re-amplifying a size-selected genomic PCR product with SHARP, rather than acting on genomic DNA directly, may help decouple the challenge of initial locus access from repeat-length fidelity, representing a readily testable next step. Future iterations of SHARP aiming for direct clinical diagnostics will likely require sequence-specific helicase loading mechanisms, macromolecular crowding agents, or optimized protein-to-template ratios to overcome the competitive genomic background.

Together, our results establish SHARP as a promising strategy for amplifying repetitive and compositionally extreme DNA sequences that are refractory to conventional PCR. By preserving primer-directed amplification while replacing global thermal denaturation with coordinated enzymatic unwinding and synthesis, SHARP provides a mechanistically distinct method to amplify difficult DNA templates. This capability could be useful for cloning, synthetic biology, sequencing library preparation, repeat locus analysis, and amplification of biologically important genomic regions that remain challenging for standard PCR workflows.

## DATA AVAILABILITY

The data underlying this article can be accessed both within the primary manuscript and its accompanying supplementary documentation.

## AUTHOR CONTRIBUTIONS

F.R. did the experimental work and prepared the manuscript.

J.K. designed and obtained preliminary data and prepared the manuscript.

T.S. and X.B. performed experiments for Figure 6.

P.J.M. performed the optical tweezer experiments.

R.M.U. provided the di-repeating 601 sequence and assisted in creating the tetra and hexa 601 sequences.

M.G. assisted with experimental design.

T.S., Y.J., A.A. and Q.Y. helped with the purification PcrA. T.A, D.B. prepared and provided Ankyrin repeat sequences.

J.M.B. guided research.

T.H. designed and directed research and prepared the manuscript.

## ACKOWLEDGEMENTS

US National Institutes of Health [grant numbers R35 GM122569 to T.H., R35 CA216778 and R37 GM071747 to J.M.B., and R01 GM068462 to D.B.] and the National Science Foundation [grant number MCB 2031094 to T.H. and J.M.B.]. T.H. is an investigator of the Howard Hughes Medical Institute (HHMI).

## MATERIAL AND METHODS

### Overview

We overexpressed and purified two principal proteins, PcrA-X M6 helicase (1 mg/mL) and *E. coli* SSB (9 mg/ml), while other components were purchased. Thermostable Inorganic PPase (NEB catalog #M0296S), Bst 3.0 DNA polymerase (8000 units/mL, NEB catalog #M0374S), ATP (100mM, ThermoFisher catalog #R0441), dNTP (10mM each, NEB catalog #N0447S). *E. coli* SSB is commercially available, and we have used it for SHARP successfully. Plasmid for PcrA-X M6 is available from Addgene (pET-11b_PcrA_M6_H93A) and recloned into a SUMO-tag plasmid for an improved expression and ease of purification (Purification details below). **Table 1** contains specific volumes of each component and stock concentrations, including our purified enzymes. However, due to batch-to-batch variation in enzyme activity, we recommend titrating enzyme volumes around the values provided in **Table 1** and monitoring SHARP activity to determine the optimal enzyme ratio for each specific batch.

**Table 1.** MasterMix components for SHARP. This table details the components and their volumes in the SHARP Master Mix for single amplification. The Reagent Mix contains ATP, dNTP, DTT, forward and reverse primers, and template DNA. The Enzyme Mix includes PcrA-X helicase, SSB (single-stranded DNA binding protein), Bst 3.0 DNA polymerase, and TIPP (thermostable inorganic pyrophosphatase). Both mixes include 2μl of 10x Buffer. The 10x Buffer contains 500mM Tris-pH7.5, 100mM (NH2)_2_SO4, 500mM KCl, 100mM MgCl2, and 20mM DTT. These buffer components support enzyme activity for the SHARP.

| Reagent Mix | Vol. (µl) | Enzyme Mix | Vol. (µl) | 10x Buffer |
| --- | --- | --- | --- | --- |
| 10x Buffer | 2.0 | 10x Buffer | 2.0 | 500 mM Tris, pH 7.5 |
| ATP (100 mM) | 2.0 | PcrA-X (1 mg/ml) | 0.5 | 100 mM (NH<sub>4</sub>)<sub>2</sub>SO<sub>4</sub> |
| dNTP (10 mM each) | 2.0 | SSB (9 mg/ml) | 1.5 | 500 mM KCl |
| Primer F (10 µM) | 2.0 | Bst 3.0 (8000 units/ml) | 0.5 | 100 mM MgCl<sub>2</sub> |
| Primer R (10 µM) | 2.0 | TPP (1000 units/ml) | 0.2 | 20 mM DTT |
| Template DNA (10 ng/µl) | 1.0 |  |  | 1 mg/ml BSA |
| ddH<sub>2</sub>O | 9.0 | ddH<sub>2</sub>O | 15.3 | pH 8.0 at 25°C |
| **Total** | **20.0** | **Total** | **20.0** |  |

### SHARP master mix

The SHARP master mix consists of two parts, a reagent mix and an enzyme mix, as shown in **Table 1**. For the reagent master mix, primer stock concentrations are at 10 μM (2 μl each for Primer F and Primer R), dNTPs (10 mM, 2 μl), ATP (100 mM, 2 μl), and 10ng of template DNA. The enzyme mix consists of SSB (9 mg/ml, 1.5 μl), PcrA-X (1 mg/mL, 0.5 μl), TIPP (100 units/ml, 0.2 μl), and Bst 3.0 (NEB catalog #M0374S) (0.5 μl). Both reagent master mix and enzyme master mix contain 2 μl of 10x buffer, and ddH2O is added to make a final volume of 20 μl each. The 10x buffer for the SHARP master mix is prepared as per the third column of the table. It contains 500 mM Tris (pH 7.5), 100 mM (NH_4_)_2_SO_4_, 500 mM KCl, 100 mM MgCl_2_, and 20 mM DTT. with a pH of 7.5 at 25°C.

### SHARP amplification protocol

For each SHARP experiment, the reagent mix and enzyme mix were prepared to a final reaction volume of 40 μL, as specified in Table 1. Reactions were incubated in a conventional PCR machine under isothermal conditions at 65°C for 45 min.

To visualize amplification products by agarose gel electrophoresis, proteins bound to the DNA were removed after the reaction, as these proteins can otherwise cause abnormal band shifts. Following amplification, SHARP products were purified using a QIAquick PCR Purification Kit. Each reaction was diluted with 5 volumes of PB buffer (Qiagen, cat. no. 19066) and loaded onto a QIAquick column. After a 1 min incubation, the column was centrifuged at 17,900 × g for 1 min, and the flow-through was discarded. The column was then washed twice with PE buffer (Qiagen, cat. no. 19065) at the same speed, with the flow-through discarded after each wash. To remove residual wash buffer, the column was placed back into the collection tube and centrifuged once more. DNA was eluted by adding 45 μL deionized water to the column, incubating for 1 min, and centrifuging to collect the purified DNA. The eluted products were analyzed on a 1% agarose gel. Each experiment was performed independently twice (n = 2), and a representative gel is shown.

### Polymerase Chain Reaction (PCR)

Conventional PCR was performed using Q5® High-Fidelity 2X Master Mix (NEB, M0492S) according to the manufacturer’s standard protocol. Annealing temperatures were determined using the NEB Tm Calculator and are listed in Table 2. PCR reactions were carried out in a 25 μL reaction volume containing 10 ng template DNA and were amplified for 30 cycles. For the mHTT amplicon, reactions contained 1 ng template DNA. The comparison was designed to evaluate SHARP against a standardized conventional PCR workflow under the conditions tested, rather than against independently optimized specialty-PCR protocols for each target. Target-specific additives, alternative polymerases, and specialized cycling programs were not systematically evaluated.

**Table 2.**
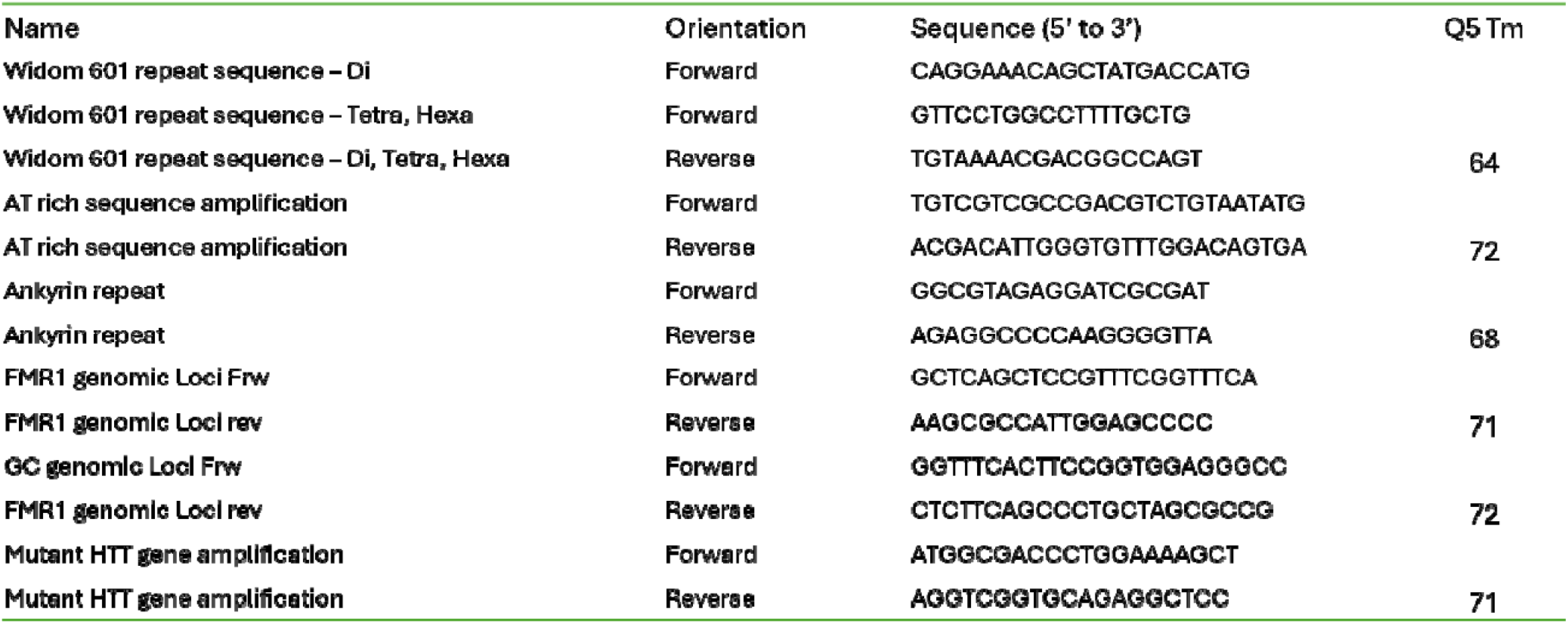
Primer pairs used in this study.

### Optical tweezer experiment protocol

Single-molecule optical tweezer experiments were performed by incubating 51 nM restriction digested SHARP amplicon and 1 nM modified DNA unzipping handles ^29^ in 1x rapid ligation buffer (Promega #C6711) with T4 DNA Ligase (NEB #M0202L) and 1 uL of 2 um anti-digoxigenin coated polystyrene particles (Spherotech #DIGP-20-2) for 10 minutes at room temperature. This reaction when then diluted in 500 uL of C-Trap buffer (1 mM EDTA pH 8, 1.5 mM MgCl2, 1 mM DTT, 0.01% BSA, 1x PBS). 2 uL of 4 um streptavidin coated polystyrene particles (Spherotech #SVP-40-5) were also diluted in 500 uL C-trap buffer. Samples were flowed in a Lumicks C-Trap flow cell in channels 1-3 in the following orientation: streptavidin-modified beads, SHARP-amplified sample incubated with anti-digoxigenin-modified beads, and C-Trap buffer. A streptavidin-coated bead and anti-digoxigenin bead were then captured in the C-Trap optical traps 1 and 2, respectively. A single DNA tether was then formed between the beads and unzipped in channel 3. Force-extension curves were fit with worm-like-chain and freely-jointed-chain models to convert extension to base pairs unzipped.

### Protein overexpression and purification

#### PcrA M6 expression and purification

PcrA M6 (H93A, C96A, N187C, C247A, L384V, L409C) was cloned into a pRSF vector with an N-terminal 12X His tag and SUMO tag using Gibson assembly. The protein was expressed in BL21(DE3) E. coli cells. Cells were grown at 37°C until reaching the density of OD 0.8, then shifted to 18°C and induced with 0.5 mM IPTG. Cells were harvested after overnight incubation at 18°C. Cell pellets were resuspended in lysis buffer (Buffer A: 50 mM HEPES pH 7.5, 1000 mM NaCl, 35 mM imidazole, 1 mM TCEP, 10% glycerol, and protease inhibitors). Cells were lysed using lysozyme and sonication, and the lysate was clarified by centrifugation. The supernatant was loaded onto a 5 mL HisTrap HP column equilibrated with Buffer A. The column was washed with 10 column volumes (CV) of Buffer A, followed by 5 CV of Buffer A containing 200 mM NaCl. The protein was then eluted directly onto a 5 mL HiTrap Heparin column using Buffer A containing 200 mM NaCl and 400 mM imidazole. Peak fractions from elution were collected and SUMO tag was cleaved off by overnight cleavage with SUMO protease. Protein was concentrated and loaded onto a HiLoad Superdex 200 pg column equilibrated with Buffer A containing 500 mM NaCl and no imidazole. Peak fractions corresponding to PcrA M6 were collected, concentrated to around 50uM, and either flash-frozen or crosslinked. For crosslinking, PcrA-M6 was treated with BMOE (bis(maleimido)ethane) crosslinker by adding a 10-20-fold molar excess of crosslinker to the protein. The mixture was incubated at room temperature for 1 hour and then further incubated on ice overnight. Excess crosslinker was removed using a PD-10 desalting column. Protein was concentrated to 1mg/ml and flash frozen in small aliquots.

#### E. coli Single Stranded binding protein (SSB)

The SSB protein was cloned into the pRSF vector without any tag using Gibson assembly. SSB was expressed and purified following a protocol adapted from Lohman et al. Briefly, the pRSF-SSB vector was transformed into E. coli BL21(DE3) cells for protein expression. The cells were grown at 37°C, induced with 0.5 mM IPTG when the optical density (OD) reached 0.8, and harvested after an additional 3 hours of growth. SSB was purified by polyethyleneimine (PEI) precipitation, followed by ammonium sulfate precipitation, and then loaded onto a HiTrap Heparin column. The protein was eluted with a gradient of NaCl. Peak fractions containing SSB were collected, concentrated to 9 mg/ml, and flash-frozen.

### Primers and templates

#### Primers

All primers used in this study were synthesized by Integrated DNA Technologies (IDT) and Azenta Life Sciences. The primer sequences are provided in **Table 2**.

#### Repetitive Widom 601 Sequences

Di-nucleosome Widom 601 sequences with 65 bp and 80 bp linker sequences were provided by Raquel Merino-Urteaga. Previous studies involving the creation of Widom 601 repetitive sequences required multiple transformation steps ^30^. Tetra- and hexa-nucleosome Widom 601 repeat sequences were generated from the di-nucleosome 601 sequences using SHARP amplification. The final construct plasmids have the following sizes: 4-65 (4,346 bp), 4-80 (4,376 bp), 6-65 (4,785 bp), and 6-80 (4,830 bp). The complete sequences of the repetitive Widom 601 amplicon are provided in the Supporting Information

#### Ankyrin Repeat Sequences

Ankyrin repeat gene sequences were previously created and reported ^31^. The sequences used in this experiment include 1, 2, 3, 5, 9, 11, and 35 repeat constructs, which were created using an amplification-ligation strategy. In our study, we observed that PCR amplification was limited to sequences with up to 4 repeats. The complete sequences of the Ankyrin repeat amplicon are provided in the Supporting Information.

#### Extreme AT-rich Sequence

The extreme AT-rich sequence plasmid template was created and provided by Ansa Biotechnologies. The sequence amplified between the primers has 95% AT content. The complete sequence of the extreme AT-rich amplicon is provided in Supporting Information.

#### Extreme GC-rich Sequence

Plasmid template for GC rich sequence was obtained from Addgene (Plasmid #63091) and genomic DNA (Coriell # NA20234)

#### mHTT Sequence

The plasmid containing the mutant HTT gene encoding exon 1 of mutant Huntington protein with a 97Q tract was a generous gift from Prof. Lu Wei at California Institute of Technology. The complete sequence of the 492bp amplicon is provided in Supporting Information.

### Nanopore sequencing and analysis

Single-molecule nanopore sequencing was performed by Plasmidsaurus (Standard Premium PCR service), which generated FASTQ files for the PCR and SHARP samples. Reads were analyzed using a custom Python (v3.12) script built on Biopython (v1.86), NumPy, pandas, and Matplotlib. Reads were filtered by mean Phred quality score (≥9) and read length (≥50 bp), retaining 8,818 of 8,842 reads (99.7%) from the PCR sample and 9,762 of 9,765 reads (99.97%) from the SHARP sample. For each expected amplicon, quality-filtered reads derived from different backbones were pooled to generate read-length histograms shown in Fig. 6B (50-bp bins, 0–1,500 bp range) and Fig. 6C (1-bp bins, 0–700 bp range; x-axis truncated at 600 bp for clarity). Enrichment of the full-length product was quantified as the fraction of reads falling within a 492 ± 10 bp window and compared between the PCR and SHARP populations. The custom Python script used for read filtering and analysis is available upon request from the corresponding author.Single-molecule sequencing was performed by Plasmidsaurus (Standard Premium PCR service). Analysis was conducted using the raw FASTQ files provided by the service.

## REFERENCES

1 Saiki, R. K. et al. Primer-directed enzymatic amplification of DNA with a thermostable DNA polymerase. Science 239, 487–491 (1988). 10.1126/science.2448875

2 Mullis, K. B. The unusual origin of the polymerase chain reaction. Sci Am 262, 56-61, 64–55 (1990). 10.1038/scientificamerican0490-56

3 Erlich, H. A., Gelfand, D. & Sninsky, J. J. Recent advances in the polymerase chain reaction. Science 252, 1643–1651 (1991). 10.1126/science.2047872

4 White, T. J., Arnheim, N. & Erlich, H. A. The polymerase chain reaction. Trends Genet 5, 185–189 (1989). 10.1016/0168-9525(89)90073-5

5 Innis, M. A., Myambo, K. B., Gelfand, D. H. & Brow, M. A. DNA sequencing with Thermus aquaticus DNA polymerase and direct sequencing of polymerase chain reaction-amplified DNA. Proc Natl Acad Sci U S A 85, 9436–9440 (1988). 10.1073/pnas.85.24.9436

6 Yang, S. & Rothman, R. E. PCR-based diagnostics for infectious diseases: uses, limitations, and future applications in acute-care settings. Lancet Infect Dis 4, 337–348 (2004). 10.1016/s1473-3099(04)01044-8

7 McDonald, C., Taylor, D. & Linacre, A. PCR in Forensic Science: A Critical Review. Genes (Basel) 15 (2024). 10.3390/genes15040438

8 Sahdev, S., Saini, S., Tiwari, P., Saxena, S. & Singh Saini, K. Amplification of GC-rich genes by following a combination strategy of primer design, enhancers and modified PCR cycle conditions. Mol Cell Probes 21, 303–307 (2007). 10.1016/j.mcp.2007.03.004

9 Hommelsheim, C. M., Frantzeskakis, L., Huang, M. & Ülker, B. PCR amplification of repetitive DNA: a limitation to genome editing technologies and many other applications. Sci Rep 4, 5052 (2014). 10.1038/srep05052

10 Mamedov, T. G. et al. A fundamental study of the PCR amplification of GC-rich DNA templates. Comput Biol Chem 32, 452–457 (2008). 10.1016/j.compbiolchem.2008.07.021

11 Chaushevska, M. et al. Get ready for short tandem repeats analysis using long reads-the challenges and the state of the art. Front Genet 16, 1610026 (2025). 10.3389/fgene.2025.1610026

12 Shinde, D., Lai, Y., Sun, F. & Arnheim, N. Taq DNA polymerase slippage mutation rates measured by PCR and quasi-likelihood analysis: (CA/GT)n and (A/T)n microsatellites. Nucleic Acids Res 31, 974–980 (2003). 10.1093/nar/gkg178

13 Walsh, P. S., Fildes, N. J. & Reynolds, R. Sequence analysis and characterization of stutter products at the tetranucleotide repeat locus vWA. Nucleic Acids Res 24, 2807–2812 (1996). 10.1093/nar/24.14.2807

14 Riet, J., Ramos, L. R. V., Lewis, R. V. & Marins, L. F. Improving the PCR protocol to amplify a repetitive DNA sequence. Genet Mol Res 16 (2017). 10.4238/gmr16039796

15 Viguera, E., Canceill, D. & Ehrlich, S. D. Replication slippage involves DNA polymerase pausing and dissociation. Embo j 20, 2587–2595 (2001). 10.1093/emboj/20.10.2587

16 Frey, U. H., Bachmann, H. S., Peters, J. & Siffert, W. PCR-amplification of GC-rich regions: ‘slowdown PCR’. Nat Protoc 3, 1312–1317 (2008). 10.1038/nprot.2008.112

17 Boán, F., Blanco, M. G., Barros, P., González, A. I. & Gómez-Márquez, J. Inhibition of DNA synthesis by K+-stabilised G-quadruplex promotes allelic preferential amplification. FEBS Lett 571, 112–118 (2004). 10.1016/j.febslet.2004.06.062

18 Zhu, X. J. et al. Guanine-rich sequences inhibit proofreading DNA polymerases. Sci Rep 6, 28769 (2016). 10.1038/srep28769

19 Dhatterwal, P., Mehrotra, S. & Mehrotra, R. Optimization of PCR conditions for amplifying an AT-rich amino acid transporter promoter sequence with high number of tandem repeats from Arabidopsis thaliana. BMC Res Notes 10, 638 (2017). 10.1186/s13104-017-2982-1

20 Richard, G. F., Kerrest, A. & Dujon, B. Comparative genomics and molecular dynamics of DNA repeats in eukaryotes. Microbiol Mol Biol Rev 72, 686–727 (2008). 10.1128/mmbr.00011-08

21 Lower, S. S., McGurk, M. P., Clark, A. G. & Barbash, D. A. Satellite DNA evolution: old ideas, new approaches. Curr Opin Genet Dev 49, 70–78 (2018). 10.1016/j.gde.2018.03.003

22 Liao, X. et al. Repetitive DNA sequence detection and its role in the human genome. Commun Biol 6, 954 (2023). 10.1038/s42003-023-05322-y

23 Mirkin, S. M. DNA structures, repeat expansions and human hereditary disorders. Curr Opin Struct Biol 16, 351–358 (2006). 10.1016/j.sbi.2006.05.004

24 Paulson, H. Repeat expansion diseases. Handb Clin Neurol 147, 105–123 (2018). 10.1016/b978-0-444-63233-3.00009-9

25 Sun, Y. J. et al. Use of mycobacterial interspersed repetitive unit-variable-number tandem repeat typing to examine genetic diversity of Mycobacterium tuberculosis in Singapore. J Clin Microbiol 42, 1986–1993 (2004). 10.1128/jcm.42.5.1986-1993.2004

26 Roychowdhury, T., Mandal, S. & Bhattacharya, A. Analysis of IS6110 insertion sites provide a glimpse into genome evolution of Mycobacterium tuberculosis. Scientific Reports 5, 12567 (2015). 10.1038/srep12567

27 Su, X. Z., Wu, Y., Sifri, C. D. & Wellems, T. E. Reduced extension temperatures required for PCR amplification of extremely A+T-rich DNA. Nucleic Acids Res 24, 1574–1575 (1996). 10.1093/nar/24.8.1574

28 Gardner, M. J. et al. Genome sequence of the human malaria parasite Plasmodium falciparum. Nature 419, 498–511 (2002). 10.1038/nature01097

29 Khaniani, M. S., Kalitsis, P., Burgess, T. & Slater, H. R. An improved Diagnostic PCR Assay for identification of Cryptic Heterozygosity for CGG Triplet Repeat Alleles in the Fragile X Gene (FMR1). Mol Cytogenet 1, 5 (2008). 10.1186/1755-8166-1-5

30 Chen, L. et al. An information-rich CGG repeat primed PCR that detects the full range of fragile X expanded alleles and minimizes the need for southern blot analysis. J Mol Diagn 12, 589–600 (2010). 10.2353/jmoldx.2010.090227

31 Green, M. R. & Sambrook, J. Nested Polymerase Chain Reaction (PCR). Cold Spring Harb Protoc 2019 (2019). 10.1101/pdb.prot095182

32 Green, M. R. & Sambrook, J. Touchdown Polymerase Chain Reaction (PCR). Cold Spring Harb Protoc 2018 (2018). 10.1101/pdb.prot095133

33 Green, M. R. & Sambrook, J. Polymerase Chain Reaction (PCR) Amplification of GC-Rich Templates. Cold Spring Harb Protoc 2019 (2019). 10.1101/pdb.prot095141

34 Chou, Q., Russell, M., Birch, D. E., Raymond, J. & Bloch, W. Prevention of pre-PCR mis-priming and primer dimerization improves low-copy-number amplifications. Nucleic Acids Res 20, 1717–1723 (1992). 10.1093/nar/20.7.1717

35 Jensen, M. A., Fukushima, M. & Davis, R. W. DMSO and betaine greatly improve amplification of GC-rich constructs in de novo synthesis. PLoS One 5, e11024 (2010). 10.1371/journal.pone.0011024

36 Notomi, T. et al. Loop-mediated isothermal amplification of DNA. Nucleic Acids Res 28, E63 (2000). 10.1093/nar/28.12.e63

37 Vincent, M., Xu, Y. & Kong, H. Helicase-dependent isothermal DNA amplification. EMBO Rep 5, 795–800 (2004). 10.1038/sj.embor.7400200

38 Gavrilov, M. et al. Engineered helicase replaces thermocycler in DNA amplification while retaining desired PCR characteristics. Nature Communications 13, 6312 (2022). 10.1038/s41467-022-34076-0

39 Segal, E. et al. A genomic code for nucleosome positioning. Nature 442, 772– 778 (2006). 10.1038/nature04979

