## Supporting Information for "Superhelicase-driven isothermal amplification of repetitive and low-complexity DNA"

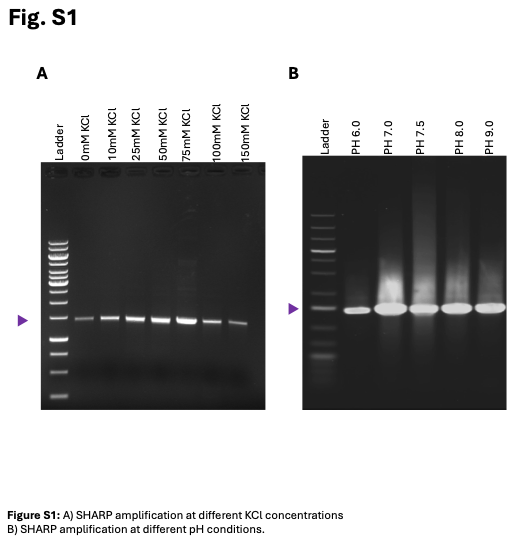

**Figure-S1**

(A) SHARP Amplification at various KCl concentrations.
(B) SHARP Amplification at various pH conditions.

**Sequences**

Sequence diversity of the amplified regions used in this study. Red indicates repeat sequences, while blue indicates variable linker sequences.

**Widom-601 2-repeats- 65bp linker** caggaaacagctatgaccatgattacgccaagcttggtaccgagctcggatccgtatacggatgtatatatcttacacgtgcctagagactagtaagtaatcctattggcggttaaaacgcggtagacagcgcgtacgtgcgtttaagcggtgctagagcttgctacgaccaattaagcggccttagcaccgggtatctccagatcctgccgagctcggatccactagtaacggccgccagtgtgctggaattcgcccttgcctgcagacaggatgtatatatcttacacgtgcctagagactagtaagtaatcctattggcggttaaaacgcggtagacagcgcgtacgtgcgtttaagcggtgctagagcttgctacgaccaattaagcggccttagcaccgggtatctccagatcgatgcaagggcctcgagcatgcatctagagggcccaattcgccctatagtgagtcgtattacaattcactggccgtcgttttaca

**Widom-601 2-repeats- 80bp linker** caggaaacagctatgaccatgattacgccaagcttggtaccgagctcggatccgtatacggatgtatatatcttacacgtgcctagagactagtaagtaatcctattggcggttaaaacgcggtagacagcgcgtacgtgcgtttaagcggtgctagagcttgctacgaccaattaagcggccttagcaccgggtatctccagatcctgacgccaagcttggtaccgagctcggatccactagtaacggccgccagtgtgctggaattcgcccttgcctgcagacaggatgtatatatcttacacgtgcctagagactagtaagtaatcctattggcggttaaaacgcggtagacagcgcgtacgtgcgtttaagcggtgctagagcttgctacgaccaattaagcggccttagcaccgggtatctccagatcgatgcaagggcctcgagcatgcatctagagggcccaattcgccctatagtgagtcgtattacaattcactggccgtcgttttaca

**Widom-601 4-repeats- 65bp linker**

gttcctggccttttgctggccttttgctcacatgtcaggaaacagctatgaccatgattacgccaagcttggtaccgagctcggatccgtatacggatgtatatatcttacacgtgcctagagactagtaagtaatcctattggcggttaaaacgcggtagacagcgcgtacgtgcgtttaagcggtgctagagcttgctacgaccaattaagcggccttagcaccgggtatctccagatcctgccgagctcggatccactagtaacggccgccagtgtgctggaattcgcccttgcctgcagacaggatgtatatatcttacacgtgcctagagactagtaagtaatcctattggcggttaaaacgcggtagacagcgcgtacgtgcgtttaagcggtgctagagcttgctacgaccaattaagcggccttagcaccgggtatctccagatcgatgcaaggccatcgtgtggacaggaaacagctatgaccatgattacgccaagcttggtaccgagctcggatccgtatacggatgtatatatcttacacgtgcctagagactagtaagtaatcctattggcggttaaaacgcggtagacagcgcgtacgtgcgtttaagcggtgctagagcttgctacgaccaattaagcggccttagcaccgggtatctccagatcctgccgagctcggatccactagtaacggccgccagtgtgctggaattcgcccttgcctgcagacaggatgtatatatcttacacgtgcctagagactagtaagtaatcctattggcggttaaaacgcggtagacagcgcgtacgtgcgtttaagcggtgctagagcttgctacgaccaattaagcggccttagcaccgggtatctccagatcgatgcaagggcctcgagcatgcatctagagggcccaattcgccctatagtgagtcgtattacaattcactggccgtcgttttaca

**Widom-601 4-repeats- 80bp linker**

gttcctggccttttgctggccttttgctcacatgtcaggaaacagctatgaccatgattacgccaagcttggtaccgagctcggatccgtatacggatgtatatatcttacacgtgcctagagactagtaagtaatcctattggcggttaaaacgcggtagacagcgcgtacgtgcgtttaagcggtgctagagcttgctacgaccaattaagcggccttagcaccgggtatctccagatcctgacgccaagcttggtaccgagctcggatccactagtaacggccgccagtgtgctggaattcgcccttgcctgcagacaggatgtatatatcttacacgtgcctagagactagtaagtaatcctattggcggttaaaacgcggtagacagcgcgtacgtgcgtttaagcggtgctagagcttgctacgaccaattaagcggccttagcaccgggtatctccagatcgatgcaaggccatcgtgtggacaggaaacagctatgaccatgattacgccaagcttggtaccgagctcggatccgtatacggatgtatatatcttacacgtgcctagagactagtaagtaatcctattggcggttaaaacgcggtagacagcgcgtacgtgcgtttaagcggtgctagagcttgctacgaccaattaagcggccttagcaccgggtatctccagatcctgacgccaagcttggtaccgagctcggatccactagtaacggccgccagtgtgctggaattcgcccttgcctgcagacaggatgtatatatcttacacgtgcctagagactagtaagtaatcctattggcggttaaaacgcggtagacagcgcgtacgtgcgtttaagcggtgctagagcttgctacgaccaattaagcggccttagcaccgggtatctccagatcgatgcaagggcctcgagcatgcatctagagggcccaattcgccctatagtgagtcgtattacaattcactggccgtcgttttaca

**Widom-601 6-repeats- 65bp linker**

gttcctggccttttgctggccttttgctcacatgtcaggaaacagctatgaccatgattacgccaagcttggtaccgagctcggatccgtatacggatgtatatatcttacacgtgcctagagactagtaagtaatcctattggcggttaaaacgcggtagacagcgcgtacgtgcgtttaagcggtgctagagcttgctacgaccaattaagcggccttagcaccgggtatctccagatcctgccgagctcggatccactagtaacggccgccagtgtgctggaattcgcccttgcctgcagacaggatgtatatatcttacacgtgcctagagactagtaagtaatcctattggcggttaaaacgcggtagacagcgcgtacgtgcgtttaagcggtgctagagcttgctacgaccaattaagcggccttagcaccgggtatctccagatcgatgcaaggccatcgtgtggacaggaaacagctatgaccatgattacgccaagcttggtaccgagctcggatccgtatacggatgtatatatcttacacgtgcctagagactagtaagtaatcctattggcggttaaaacgcggtagacagcgcgtacgtgcgtttaagcggtgctagagcttgctacgaccaattaagcggccttagcaccgggtatctccagatcctgccgagctcggatccactagtaacggccgccagtgtgctggaattcgcccttgcctgcagacaggatgtatatatcttacacgtgcctagagactagtaagtaatcctattggcggttaaaacgcggtagacagcgcgtacgtgcgtttaagcggtgctagagcttgctacgaccaattaagcggccttagcaccgggtatctccagatcgatgcaaggcacagcgtgcacaggaaacagctatgaccatgattacgccaagcttggtaccgagctcggatccgtatacggatgtatatatcttacacgtgcctagagactagtaagtaatcctattggcggttaaaacgcggtagacagcgcgtacgtgcgtttaagcggtgctagagcttgctacgaccaattaagcggccttagcaccgggtatctccagatcctgccgagctcggatccactagtaacggccgccagtgtgctggaattcgcccttgcctgcagacaggatgtatatatcttacacgtgcctagagactagtaagtaatcctattggcggttaaaacgcggtagacagcgcgtacgtgcgtttaagcggtgctagagcttgctacgaccaattaagcggccttagcaccgggtatctccagatcgatgcaagggcctcgagcatgcatctagagggcccaattcgccctatagtgagtcgtattacaattcactggccgtcgttttaca

**Widom-601 6-repeats- 80bp linker**

gttcctggccttttgctggccttttgctcacatgtcaggaaacagctatgaccatgattacgccaagcttggtaccgagctcggatccgtatacggatgtatatatcttacacgtgcctagagactagtaagtaatcctattggcggttaaaacgcggtagacagcgcgtacgtgcgtttaagcggtgctagagcttgctacgaccaattaagcggccttagcaccgggtatctccagatcctgacgccaagcttggtaccgagctcggatccactagtaacggccgccagtgtgctggaattcgcccttgcctgcagacaggatgtatatatcttacacgtgcctagagactagtaagtaatcctattggcggttaaaacgcggtagacagcgcgtacgtgcgtttaagcggtgctagagcttgctacgaccaattaagcggccttagcaccgggtatctccagatcgatgcaaggccatcgtgtggacaggaaacagctatgaccatgattacgccaagcttggtaccgagctcggatccgtatacggatgtatatatcttacacgtgcctagagactagtaagtaatcctattggcggttaaaacgcggtagacagcgcgtacgtgcgtttaagcggtgctagagcttgctacgaccaattaagcggccttagcaccgggtatctccagatcctgacgccaagcttggtaccgagctcggatccactagtaacggccgccagtgtgctggaattcgcccttgcctgcagacaggatgtatatatcttacacgtgcctagagactagtaagtaatcctattggcggttaaaacgcggtagacagcgcgtacgtgcgtttaagcggtgctagagcttgctacgaccaattaagcggccttagcaccgggtatctccagatcgatgcaaggcacagcgtgcacaggaaacagctatgaccatgattacgccaagcttggtaccgagctcggatccgtatacggatgtatatatcttacacgtgcctagagactagtaagtaatcctattggcggttaaaacgcggtagacagcgcgtacgtgcgtttaagcggtgctagagcttgctacgaccaattaagcggccttagcaccgggtatctccagatcctgacgccaagcttggtaccgagctcggatccactagtaacggccgccagtgtgctggaattcgcccttgcctgcagacaggatgtatatatcttacacgtgcctagagactagtaagtaatcctattggcggttaaaacgcggtagacagcgcgtacgtgcgtttaagcggtgctagagcttgctacgaccaattaagcggccttagcaccgggtatctccagatcgatgcaagggcctcgagcatgcatctagagggcccaattcgccctatagtgagtcgtattacaattcactggccgtcgttttaca

**Ankyrin-1repeat**

ggcgtagaggatcgcgatctcgatcccgcgaaattaatacgactcactataggggaattgtgagcggataacaattcccctctagaaataattttgtttaactttaagaaggagatataccatgggccatatgtggggatccaaagatggcaacaccccgctgcataacgcggcgaaaaacggccatgcggaagaagtgaaaaaactgctgtccaaaggcgcggatgtgaacgcgagatcttggggcagcagccatcatcatcatcatcattagctgagcaataactagcataaccccttggggcctct

**Ankyrin-2repeat**

ggcgtagaggatcgcgatctcgatcccgcgaaattaatacgactcactataggggaattgtgagcggataacaattcccctctagaaataattttgtttaactttaagaaggagatataccatgggccatatgtggggatccaaagatggcaacaccccgctgcataacgcggcgaaaaacggccatgcggaagaagtgaaaaaactgctgtccaaaggcgcggatgtgaacgcgagatccaaagatggcaacaccccgctgcatctggcggcgaaaaacggccatgcggaaatcgtgaaactgctgctggcgaaaggcgcggatgtgaacgcgagatcttggggcagcagccatcatcatcatcatcattagctgagcaataactagcataaccccttggggcctct

**Ankyrin-3repeat**

ggcgtagaggatcgcgatctcgatcccgcgaaattaatacgactcactataggggaattgtgagcggataacaattcccctctagaaataattttgtttaactttaagaaggagatataccatgggccatatgtggggatccaaagatggcaacaccccgctgcataacgcggcgaaaaacggccatgcggaagaagtgaaaaaactgctgtccaaaggcgcggatgtgaacgcgagatccaaagatggcaacaccccgctgcatctggcggcgaaaaacggccatgcggaaatcgtgaaactgctgctggcgaaaggcgcggatgtgaacgcgagatccaaagatggcaacaccccgctgcatctggcggcgaaaaacggccatgcggaaatcgtgaaactgctgctggcgaaaggcgcggatgtgaacgcgagatcttggggcagcagccatcatcatcatcatcattagctgagcaataactagcataaccccttggggcctct

**Ankyrin-5repeat**

ggcgtagaggatcgcgatctcgatcccgcgaaattaatacgactcactataggggaattgtgagcggataacaattcccctctagaaataattttgtttaactttaagaaggagatataccatgggccatatgtggggatccaaagatggcaacaccccgctgcataacgcggcgaaaaacggccatgcggaagaagtgaaaaaactgctgtccaaaggcgcggatgtgaacgcgagatccaaagatggcaacaccccgctgcatctggcggcgaaaaacggccatgcggaaatcgtgaaactgctgctggcgaaaggcgcggatgtgaacgcgagatccaaagatggcaacaccccgctgcatctggcggcgaaaaacggccatgcggaaatcgtgaaactgctgctggcgaaaggcgcggatgtgaacgcgagatccaaagatggcaacaccccgctgcatctggcggcgaaaaacggccatgcggaaatcgtgaaactgctgctggcgaaaggcgcggatgtgaacgcgagatccaaagatggcaacaccccgctgcatctggcggcgaaaaacggccatgcggaaatcgtgaaactgctgctggcgaaaggcgcggatgtgaacgcgagatcttggggcagcagccatcatcatcatcatcattagctgagcaataactagcataaccccttggggcctct

**Ankyrin-9repeat**

ggcgtagaggatcgcgatctcgatcccgcgaaattaatacgactcactataggggaattgtgagcggataacaattcccctctagaaataattttgtttaactttaagaaggagatataccatgggccatatgtggggatccaaagatggcaacaccccgctgcataacgcggcgaaaaacggccatgcggaagaagtgaaaaaactgctgtccaaaggcgcggatgtgaacgcgagatccaaagatggcaacaccccgctgcatctggcggcgaaaaacggccatgcggaaatcgtgaaactgctgctggcgaaaggcgcggatgtgaacgcgagatccaaagatggcaacaccccgctgcatctggcggcgaaaaacggccatgcggaaatcgtgaaactgctgctggcgaaaggcgcggatgtgacgcgagatccaaagatggcaacaccccgctgcatctggcggcgaaaaacggccatgcggaaatcgtgaaactgctgctggcgaaaggcgcggatgtgaacgcgagatccaaagatggcaacaccccgctgcatctggcggcgaaaaacggccatgcggaaatcgtgaaactgctgctggcgaaaggcgcggatgtgaacgcgagatccaaagatggcaacaccccgctgcatctggcggcgaaaaacggccatgcggaaatcgtgaaactgctgctggcgaaaggcgcggatgtgaacgcgagatccaaagatggcaacaccccgctgcatctggcggcgaaaaacggccatgcggaaatcgtgaaactgctgctggcgaaaggcgcggatgtgaacgcgagatccaaagatggcaacaccccgctgcatctggcggcgaaaaacggccatgcggaaatcgtgaaactgctgctggcgaaaggcgcggatgtgaacgcgagatccaaagatggcaacaccccgctgcatctggcggcgaaaaacggccatgcggaaatcgtgaaactgctgctggcgaaaggcgcggatgtgaacgcgagatcttggggcagcagccatcatcatcatcatcattagctgagcaataactagcataaccccttggggcctct

**Ankyrin-11repeat**

ggcgtagaggatcgcgatctcgatcccgcgaaattaatacgactcactataggggaattgtgagcggataacaattcccctctagaaataattttgtttaactttaagaaggagatataccatgggccatatgtggggatccaaagatggcaacaccccgctgcataacgcggcgaaaaacggccatgcggaagaagtgaaaaaactgctgtccaaaggcgcggatgtgaacgcgagatccaaagatggcaacaccccgctgcatctggcggcgaaaaacggccatgcggaaatcgtgaaactgctgctggcgaaaggcgcggatgtgaacgcgagatccaaagatggcaacaccccgctgcatctggcggcgaaaaacggccatgcggaaatcgtgaaactgctgctggcgaaaggcgcggatgtgaacgcgagatccaaagatggcaacaccccgctgcatctggcggcgaaaaacggccatgcggaaatcgtgaaactgctgctggcgaaaggcgcggatgtgaacgcgagatccaaagatggcaacaccccgctgcatctggcggcgaaaaacggccatgcggaaatcgtgaaactgctgctggcgaaaggcgcggatgtgaacgcgagatccaaagatggcaacaccccgctgcatctggcggcgaaaaacggccatgcggaaatcgtgaaactgctgctggcgaaaggcgcggatgtgaacgcgagatccaaagatggcaacaccccgctgcatctggcggcgaaaaacggccatgcggaaatcgtgaaactgctgctggcgaaaggcgcggatgtgaacgcgagatccaaagatggcaacaccccgctgcatctggcggcgaaaaacggccatgcggaaatcgtgaaactgctgctggcgaaaggcgcggatgtgaacgcgagatccaaagatggcaacaccccgctgcatctggcggcgaaaaacggccatgcggaaatcgtgaaactgctgctggcgaaaggcgcggatgtgaacgcgagatccaaagatggcaacaccccgctgcatctggcggcgaaaaacggccatgcggaaatcgtgaaactgctgctggcgaaaggcgcggatgtgaacgcgagatccaaagatggcaacaccccgctgcatctggcggcgaaaaacggccatgcggaaatcgtgaaactgctgctggcgaaaggcgcggatgtgaacgcgagatccaaagatggcaacaccccgctgcatctggcggcgaaaaacggccatgcggaaatcgtgaaactgctgctggcgaaaggcgcggatgtgaacgcgagatcttggggcagcagccatcatcatcatcatcattagctgagcaataactagcataaccccttggggcctct

**Ankyririn-35repeat**

ggcgtagaggatcgcgatctcgatcccgcgaaattaatacgactcactataggggaattgtgagcggataacaattcccctctagaaataattttgtttaactttaagaaggagatataccatgggccatatgtggggatccaaagatggcaacaccccgctgcataacgcggcgaaaaacggccatgcggaagaagtgaaaaaactgctgtccaaaggcgcggatgtgaacgcgagatccaaagatggcaacaccccgctgcatctggcggcgaaaaacggccatgcggaaatcgtgaaactgctgctggcgaaaggcgcggatgtgaacgcgagatccaaagatggcaacaccccgctgcatctggcggcgaaaaacggccatgcggaaatcgtgaaactgctgctggcgaaaggcgcggatgtgaaacgcgagatccaaagatggcaacaccccgctgcatctggcggcgaaaaacggccatgcggaaatcgtgaaactgctgctggcgaaaggcgcggatgtgaacgcgagatccaaagatggcaacaccccgctgcatctggcggcgaaaaacggccatgcggaaatcgtgaaactgctgctggcgaaaggcgcggatgtgaaacgcgagatccaaagatggcaacaccccgctgcatctggcggcgaaaaacggccatgcggaaatcgtgaaactgctgctggcgaaaggcgcggatgtgaacgcgagatccaaagatggcaacaccccgctgcatctggcggcgaaaaacggccatgcggaaatcgtgaaactgctgctggcgaaaggcgcggatgtgaaacgcgagatccaaagatggcaacaccccgctgcatctggcggcgaaaaacggccatgcggaaatcgtgaaactgctgctggcgaaaggcgcggatgtgaacgcgagatccaaagatggcaacaccccgctgcatctggcggcgaaaaacggccatgcggaaatcgtgaaactgctgctggcgaaaggcgcggatgtgaaacgcgagatccaaagatggcaacaccccgctgcatctggcggcgaaaaacggccatgcggaaatcgtgaaactgctgctggcgaaaggcgcggatgtgaacgcgagatccaaagatggcaacaccccgctgcatctggcggcgaaaaacggccatgcggaaatcgtgaaactgctgctggcgaaaggcgcggatgtgaaacgcgagatccaaagatggcaacaccccgctgcatctggcggcgaaaaacggccatgcggaaatcgtgaaactgctgctggcgaaaggcgcggatgtgaacgcgagatccaaagatggcaacaccccgctgcatctggcggcgaaaaacggccatgcggaaatcgtgaaactgctgctggcgaaaggcgcggatgtgaacgcgagatccaaagatggcaacaccccgctgcatctggcggcgaaaaacggccatgcggaaatcgtgaaactgctgctggcgaaaggcgcggatgtgaacgcgagatccaaagatggcaacaccccgctgcatctggcggcgaaaaacggccatgcggaaatcgtgaaactgctgctggcgaaaggcgcggatgtgaacgcgagatccaaagatggcaacaccccgctgcatctggcggcgaaaaacggccatgcggaaatcgtgaaactgctgctggcgaaaggcgcggatgtgaacgcgagatccaaagatggcaacaccccgctgcatctggcggcgaaaaacggccatgcggaaatcgtgaaactgctgctggcgaaaggcgcggatgtgaacgcgagatccaaagatggcaacaccccgctgcatctggcggcgaaaaacggccatgcggaaatcgtgaaactgctgctggcgaaaggcgcggatgtgaacgcgagatccaaagatggcaacaccccgctgcatctggcggcgaaaaacggccatgcggaaatcgtgaaactgctgctggcgaaaggcgcggatgtgaacgcgagatccaaagatggcaacaccccgctgcatctggcggcgaaaaacggccatgcggaaatcgtgaaactgctgctggcgaaaggcgcggatgtgaacgcgagatccaaagatggcaacaccccgctgcatctggcggcgaaaaacggccatgcggaaatcgtgaaactgctgctggcgaaaggcgcggatgtgaacgcgagatccaaagatggcaacaccccgctgcatctggcggcgaaaaacggccatgcggaaatcgtgaaactgctgctggcgaaaggcgcggatgtgaacgcgagatccaaagatggcaacaccccgctgcatctggcggcgaaaaacggccatgcggaaatcgtgaaactgctgctggcgaaaggcgcggatgtgaacgcgagatccaaagatggcaacaccccgctgcatctggcggcgaaaaacggccatgcggaaatcgtgaaactgctgctggcgaaaggcgcggatgtgaacgcgagatccaaagatggcaacaccccgctgcatctggcggcgaaaaacggccatgcggaaatcgtgaaactgctgctggcgaaaggcgcggatgtgaacgcgagatccaaagatggcaacaccccgctgcatctggcggcgaaaaacggccatgcggaaatcgtgaaactgctgctggcgaaaggcgcggatgtgaacgcgagatccaaagatggcaacaccccgctgcatctggcggcgaaaaacggccatgcggaaatcgtgaaactgctgctggcgaaaggcgcggatgtgaacgcgagatccaaagatggcaacaccccgctgcatctggcggcgaaaaacggccatgcggaaatcgtgaaactgctgctggcgaaaggcgcggatgtgaacgcgagatccaaagatggcaacaccccgctgcatctggcggcgaaaaacggccatgcggaaatcgtgaaactgctgctggcgaaaggcgcggatgtgaacgcgagatccaaagatggcaacaccccgctgcatctggcggcgaaaaacggccatgcggaaatcgtgaaactgctgctggcgaaaggcgcggatgtgaacgcgagatccaaagatggcaacaccccgctgcatctggcggcgaaaaacggccatgcggaaatcgtgaaactgctgctggcgaaaggcgcggatgtgaacgcgagatccaaagatggcaacaccccgctgcatctggcggcgaaaaacggccatgcggaaatcgtgaaactgctgctggcgaaaggcgcggatgtgaacgcgagatccaaagatggcaacaccccgctgcatctggcggcgaaaaacggccatgcggaaatcgtgaaactgctgctggcgaaaggcgcggatgtgaacgcgagatccaaagatggcaacaccccgctgcatctggcggcgaaaaacggccatgcggaaatcgtgaaactgctgctggcgaaaggcgcggatgtgaacgcgagatcttggggcagcagccatcatcatcatcatcattagctgagcaataactagcataaccccttggggcctct

**AT-Rich seq-95%**

tggaattatcactgcgtgcggcatgaagattataaattaatattaatatttatatataaaattatttaatataattataaatatttttaaattatatttttataaattattataaaatttttaattttaattattatattattttaatattaaatataaaaataaaaattatttaattaaatatttataaattattttataatattatatatataaattaatattattaataatttaaaatattatataattaaatatatttacccggtccaatttaatgttgatactgtaagggtggatcctctataataatagatattttatattttaaattattaataatattaatccatatatataatattataaaataatttataaatatttaattaaataattttttttaaattatatttttataaattattataaaatttttaattttaattattatattattttaatattaaatataaaaataatatttataattatattaaataattttatatataaatattaatattaatttataatcttcatgccgcacgtattagtcttcatacgaacctcaa

**FMR1 Genomic loci-83% GC (CORIELL # NA20234)**

GCTCAGCTCCGTTTCGGTTTCACTTCCGGTGGAGGGCCGCCTCTGAGCGGGCGGCGGGCCGACGGCGAGCGCGGGCGGCGGCGGTGACGGAGGCGCCGCTGCCAGGGGGCGTGCGGCAGCGCGGCGGCGGCGGCGGCGGCGGCGGCGGCGGAGGCGGCGGCGGCGGCGGCGGCGGCGGCGGCTGGGCCTCGAGCGCCCGCAGCCCACCTCTCGGGGGCGGGCTCCCGGCGCTAGCAGGGCTGAAGAGAAGATGGAGGAGCTGGTGGTGGAAGTGCGGGGCTCCAATGGCGCTT

**CGG repeat plasmid-95% GC pMAM-CGG 99x FMR1 (Addgene Plasmid #63091)**

ggtttcacttccggtggagggccgcctctgagcgggcggcgggccgacggcgagcgcgggcggcggcggtgacggaggcgccgctgccagggggcgtgcggcagcgcggcggcggcggcggcggcggcggcggcggcggcggcggcggaggcggcggcggcggcggcggcggcggcggcggcggcggcggcggcggcggcggcggcggcggcggcggcggcggcggcggcggcggcggcggcggcggcggcggcggcggcggcggcggcggcggcggcggcggcggcggcggcggcggcggcggcggcggcggcggcggcggcggcggcggcggcggcggcggcggcggcggcggcggcggcggcggcggcggcggcggcggcggcggcggcggcggcggcggctgggcctcgagcgcccgcagcccacctctcgggggcgggctcccggcgctagcagggctgaagag

**mHTT Sequence**

The plasmid containing the mutant HTT gene encoding exon 1 of mutant Huntington protein with a 97Q tract was a generous gift from Prof. Lu Wei at California Institute of Technology. The sequence of the 492 bp amplicon is shown below, with the region encoding the 97Q tract highlighted in bold.

ATGGCGACCCTGGAAAAGCTGATGAAGGCCTTCGAGTCCCTCAAAAGCTTC

**CAA CAG CAG CAA CAG CAA CAA CAG CAG CAA CAG CAA CAA CAG CAG CAA CAG CAA CAA CAG CAG CAA CAG CAA CAA CAG CAG CAA CAG CAA CAA CAG CAG CAA CAG CAA CAA CAG CAG CAA CAG CAA CAA CAG CAG CAA CAG CAA CAA CAG CAG CAA CAG CAA CAA CAG CAG CAA CAG CAA CAA CAG CAG CAA CAG CAA CAA CAG CAG CAA CAG CAA CAA CAG CAG CAA CAG CAA CAA CAG CAG CAA CAG CAA CAA CAG CAG CAA CAG CAA CAA CAG CAG CAA CAG CAA CAA**

CCGCCACCACCTCCCCCTCCACCCCCACCTCCTCAACTTCCTCAACCTCCTCCACAGGCACAGCCTCTGCTGCCTCAGCCACAACCTCCTCCACCTCCACCTCCACCTCCTCCAGGCCCAGCTGTGGCTGAGGAGCCTCTGCACCGACCT
